# Taxa-driven functional shifts associated with stormflow in an urban stream microbial community

**DOI:** 10.1101/300699

**Authors:** Adit Chaudhary, Imrose Kauser, Anirban Ray, Rachel Poretsky

## Abstract

Urban streams are susceptible to stormwater and sewage inputs that can impact their ecological health and water quality. Microbial communities in streams play important functional roles and their composition and metabolic potential can help assess ecological state and water quality. Although these environments are highly heterogenous, little is known about the influence of isolated perturbations, such as those resulting from rain events on urban stream microbiota. Here, we examined the microbial community composition and diversity in an urban stream during dry and wet weather conditions with both 16S rRNA gene sequencing across multiple years and shotgun metagenomics to more deeply analyze a single stormflow event. Metagenomics was used to assess population-level dynamics as well as shifts in the microbial community taxonomic profile and functional potential before and after a substantial rainfall. Results demonstrated general trends present in the stream under stormflow vs. baseflow conditions across years and seasons and also highlighted the significant influence of increased effluent flow following rain in shifting the stream microbial community from abundant freshwater taxa to those more associated with urban/anthropogenic settings. Shifts in the taxonomic composition were also linked to changes in functional gene content, particularly for transmembrane transport and organic substance biosynthesis. We also observed an increase in relative abundance of genes encoding degradation of organic pollutants and antibiotic resistance after rain. Overall, this study provided evidence of stormflow impacts on an urban stream microbiome from an environmental and public health perspective.

**Importance:** Urban streams in various parts of the world are facing increased anthropogenic pressure on their water quality, and stormflow events represent one such source of complex physical, chemical and biological perturbations. Microorganisms are important components of these streams from both ecological and public-health perspectives, and analyzing the effect of such perturbations on the stream microbial community can help improve current knowledge on the impact such chronic disturbances can have on these water resources. This study examines microbial community dynamics during rain-induced stormflow conditions in an urban stream of the Chicago Area Waterway System. Additionally, using shotgun metagenomics we identified significant shifts in the microbial community composition and functional gene content following a high rainfall event, with potential environment and public health implications. Previous work in this area has been limited to specific genes/organisms or has not assessed immediate stormflow impact.

## Introduction

Streams and rivers are important freshwater resources, used for recreation, agriculture, domestic water sources and industrial purposes. By storing, processing, and transporting terrestrially derived nutrients and organic matter, rivers play an important ecological role in linking biogeochemical cycles between terrestrial and aquatic ecosystems (1). Over the last century, many streams and rivers have witnessed rapid urbanization and anthropogenic development of their drainage basins, which has exposed them to frequent external inputs in the form of wastewater treatment plant (WWTP) effluent, industrial discharge, and sewer/stormwater overflows. These inputs often impact stream hydrological, physicochemical, and biological characteristics (2). For streams and rivers that serve as wastewater and/or stormwater outfall sites, rain-induced stormflow events are especially influential, as they often lead to an increased influx of WWTP effluent and unregulated waste via combined sewer overflows (CSOs) (3, 4). These perturbations bring in nutrients, a variety of microorganisms including pathogens, and chemical pollutants such as steroid hormones that impact water quality, biodiversity, and ecosystem health (2, 3, 5, 6).

Because urban aquatic streams are typically highly variable systems that are regularly subject to anthropogenic inputs, it is unclear how much isolated perturbations such as rainfall and associated increases in stormflow might influence the water column microbial community, even in the short-term. Studies investigating urban river microbiota using genetic markers for fecal bacteria or 16S rRNA gene-based microbial community surveys have shown the presence of human fecal contamination, ‘urban signature’ bacteria and changes in community composition in streams and rivers impacted by WWTP effluent, stormwater, and CSOs (7–11). Moreover, others have documented the possible influx of antibiotic resistance bacteria and pathogens from WWTP effluent (12, 13) and stormwater events (6, 14) into urban environments, further signifying the importance of evaluating the persistence of these organisms and their impact on the riverine microbiome from a public health perspective. While these studies provide valuable information about the effects of stormflow events on urban stream microbial content they are limited to specific taxonomic and pollutant marker genes. Recent whole genome shotgun (WGS) metagenomics-based approaches have explored community composition and functional dynamics in urban impacted streams (15, 16), although a direct effect of stormflow on microbial dynamics remains less explored. A robust evaluation of the impacts of such isolated and shortterm perturbations is critical for making predictions about the public health and possible longer-term ecological implications.

In this study, we used both 16S rRNA gene amplicon and shotgun metagenomics to deeply analyze the water column microbial community during baseflow and stormflow conditions in the North Shore Channel (NSC) stream, a section of the highly urbanized Chicago Area Waterway System (CAWS) (Fig. S1). We focused on a site downstream of a WWTP and numerous CSO outflow points using 16S rRNA gene amplicon sequencing of samples from both baseflow and stormflow over the course of multiple seasons and years. Additionally, samples obtained immediately before and shortly (<24 hr) after a single rain event at the same site provided an opportunity for a deep analysis of short-term variability in the taxonomic and functional composition of the water-column microbiome using WGS metagenomics. Coupled with the 16S rRNA data from multiple samples, we were able to link some of these changes to stormflow conditions. We identified notable rainfall-induced changes in the stream microbial taxonomic and functional profiles, driven by shifts in the relative abundance of a few abundant microbial groups such as Actinobacteria and *Legionella* that could be functionally linked to the processes of transmembrane transport and organic substance biosynthesis. We also observed an increase in genes associated with antibiotic resistance and biodegradation of known wastewater pollutants following rain. Although our deep metagenomics-based analysis is centered around a single event, our findings provides a window into the variability and short-term changes in an urban freshwater system and sets the groundwork for making predictions about possible ecosystem level and public health related impacts of rainfall events on these systems. Overall, our results show that rain-associated WWTP effluent flow and perhaps CSOs impact the stream microbiome composition and functional potential, with the introduction of exogenous organisms to the system being a significant driver of the observed change.

## Results and Discussion

### Impact of rainfall on North Shore Channel (NSC) microbial community composition

Rainfall can impact urban waterways by increasing effluent flow from WWTPs or causing combined sewer overflow events (CSOs) at outflow points along streams (4). The NSC site that we investigated has a WWTP (O’Brien Water Reclamation Plant) and several CSO outfall sites within a few km upstream (Fig. S1) and often experiences increased flow from both following rainfall, including the two rain events reported in this study (http://www.mwrd.org/irj/portal/anonymous/overview)(Fig. S2). Sequences from 16S rRNA gene amplicons at five distinct times between 2013-2015 representing both summer and fall and stream baseflow (dry weather; three samples) and stormflow (<24 hours after rain; two samples) (additional details are in Table S1) revealed both a seasonal/temporal and rainfall-associated clustering of the samples at the OTU level (PCoA, Bray-Curtis metric) (Fig. 1A). In particular, the separate clustering of stormflow and baseflow samples along the Prinicipal Axis 2 highlights the strong influence of rain on the microbial community composition across different seasons. Such changes might result from either a direct influx of allochthonous microbes or a shift in the resident microbial community in response to altered chemical conditions following rain, although none of the measured physicochemical parameters showed a statistically significant difference between stormflow and baseflow conditions (p > 0.05, Welch’s t-test; Table S1). In addition to shifts in community composition, microbial diversity based on OTU richness and Good’s coverage was slightly higher in the stormflow samples than the baseflow samples (Table S2), although the differences were not significant (p > 0.05, Welch’s t-test).

**Figure 1:**
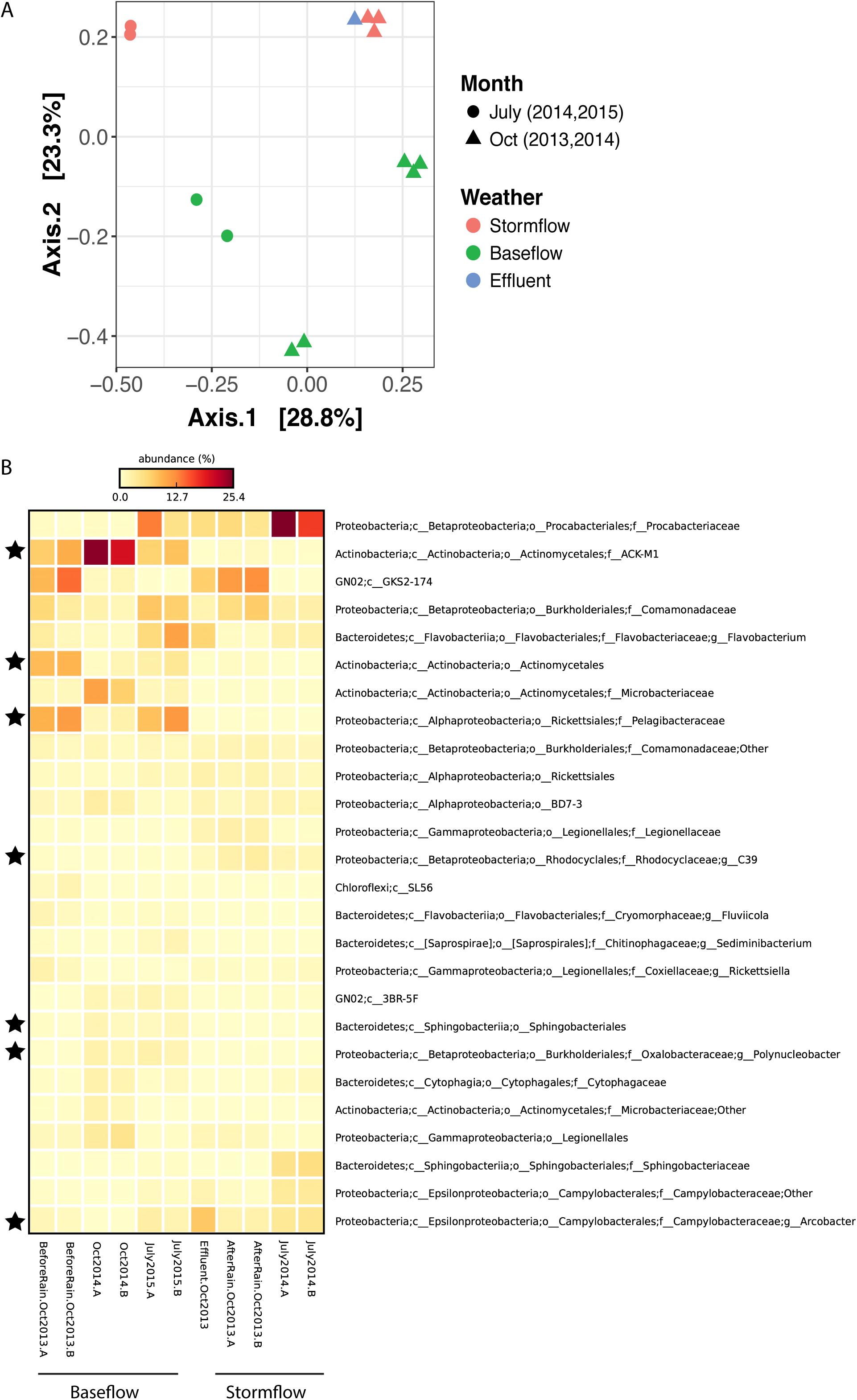
**(A)** Principal coordinate analysis (PCoA) (Bray-Curtis metric) of OTU-based microbial community diversity for North Shore Channel (NSC) water and WWTP effluent. Samples were obtained during either baseflow or stormflow conditions between 2013-2015 in summer (July) and fall (October) seasons. Each NSC time point is represented on the PCoA by biological duplicates, except for Oct-2013 stormflow and baseflow samples that also have sequencing duplicates for one of their bio-samples. **(B)** Heat map representing the relative abundance (percent of total 16S rRNA gene sequences) of dominant bacterial taxa classified till the lowest possible level (up-to genus) for the NSC and effluent samples. Taxa highlighted with a star symbol represent bacterial groups with significantly different relative abundance (p < 0.05, Welch’s t-test) between the stormflow and baseflow samples of NSC. Two biological replicates marked as A and B represent each NSC time point, and the average value of these replicates per time point was used in Welch’s t-test between the two groups (stormflow and baseflow).

To analyze shifts in the microbial community across all stormflow vs. baseflow samples, OTUs were clustered at various hierarchical taxonomic levels. There was a difference in genus-based community composition between the stormflow and baseflow samples as per ANOSIM (Bray-Curtis metric, R^2^ = 0.5, p = 0.1). Genus-level comparisons of microbial community composition revealed a significantly lower abundance of unknown genera within groups *Pelagibacteraceae*, ACK-M1 and *Actinomycetales* and a significantly higher abundance of *Arcobacter* and unknown genera within the family *Rhodocyclaceae* during stormflow as compared to baseflow (p < 0.05, Welch’s t-test) (Fig. 1B). The ACK-M1 family of Actinobacteria and *Pelagibacteraceae* are common freshwater organisms that do not favor nutrient rich conditions (17, 18) while genera within *Rhodocycleae* are Betaproteobacteria known to take advantage of nutrient/substrate-rich conditions, likely due to higher growth rates (17). *Rhodocycleae* has previously been associated with urban streams and was reported to be abundant in impacted Milwaukee waterways (19). Similarly, *Arcobacter* has often been associated with sewage and WWTP effluent (8, 9, 20). The increase in the relative abundance of these organisms in the NSC following rainfall could be due to point source inputs from the increased effluent flow and/or CSOs and was analyzed in more detail with shotgun metagenomics (see below).

Overall, the rain-associated changes in the microbial community composition appeared to be directly related to increased effluent; the after rain community OTUs were more similar to those in the WWTP effluent than to the before rain community (Fig. 1A). This could be linked to a few taxa, such as unknown genera within families *Procabacteriaceae* and *Legionellacaea* as well as the genus *Arcobacter*, which were abundant in the effluent and increased in the stream post-rain (Fig. 1B).

### Metagenomics-based microbial community composition before and after rain in North Shore Channel

The overall trends from the 16S rRNA gene-based analysis across seasons and years warranted a whole community metagenomic analysis of more temporally resolved samples clustered around a large rainfall event. Metagenomes with 4.06-16.21 million reads per library were obtained (Table S3) from the same NSC site discussed above (Fig. S1) before and <24 h after a heavy rainfall that followed a dry period in October 2013 (Fig. S2). These were used to comprehensively identify short-term changes in the microbial taxonomic profile after the rain. The rain resulted in increased WWTP effluent flow into the stream for ~24 h following precipitation, from <200 MGD to >300 MGD, and several CSO events at at-least three outfall locations upstream of sampled site within 10 h of rain (http://www.mwrd.org/irj/portal/anonymous/overview)(Fig. S2). Community coverage estimates using read redundancy (21) showed that the before rain metagenomes captured between 50-60% of the community and the after rain libraries captured approximately 40% (Fig. S3), indicating only a nominal increase in diversity after rainfall; a small increase in community OTU richness after rain was also observed with the 16S rRNA gene amplicon data (Table S2). Furthermore, the concentration of microbial cells in the before and after rain samples were similar: 1.39×10^6^ and 1.25×10^6^ cells/ml, respectively. Previous studies have reported conflicting responses of microbial community diversity to urban inputs, with some showing an increase (20) and others a decrease (15, 23) relative to less impacted conditions/systems. This may be due to differing base conditions; the NSC is characterized by significant urban effluent flow even in the absence of rain. While Lake Michigan provides the primary freshwater input, about 70% of the annual flow through the CAWS is contributed by the treated effluent discharge from WWTPs in the city (24) during both baseflow and stormflow conditions. Our results do not show a strong pattern of change in microbial community diversity/richness during stormflow in NSC, perhaps because of the variable nature of urban stream microbial communities or due to the small size of the this study. However, we hypothesize based on our results that individual rain events might not significantly impact microbial diversity in this system.

Despite overall similarities in microbial diversity and cell counts, numerous taxonomic differences were seen following rain, indicating that these changes likely reflect actual changes in microbial populations. The microbial communities pre- and post-rainfall determined both from 16S rRNA genes and by assigning taxa to assembled metagenomic contigs showed overall concordance, however we focused on the assembled contigs for a high-resolution, population-level characterization of the community and to evaluate possible links between taxonomic and functional changes in the microbiome (24). About ~67% of the large (>500bp) contigs used by MyTaxa were classifiable at phylum level, ~35% at genus level, and 24% at species level. At the phylum level (Proteobacteria subdivided into classes), several individual taxa showed significantly different relative abundances after rain with large effect sizes (Fig. 2A). Actinobacteria and Bacteroidetes significantly decreased in relative abundance after rain whereas Gammaproteobacteria, Betaproteobacteria and Chlamydiae significantly increased (p <0.05, t-test, false discovery rate corrected) (Fig. 2A). Similarity Percentage analysis (SIMPER, 26) revealed that Actinobacteria, Gammaproteobacteria and unclassified Proteobacteria contributed the most (35%, 14% and 21%, respectively) to the differences in community composition between the before and after rain samples at the phylum level. At the genus level, the decrease in relative abundance of Innominate (unclassified at genus level) Actinobacteria, *Candidatus Pelagibacter* and *Streptomyces* as well as the increase in relative abundance of *Legionella* and *Rickettsia*-affiliated sequences after rain contributed to the major change (>50%) in community composition (Fig. 2B). *Francisella*, *Nitrospira*, *Chlamydia* and *Pseudomonas* were other major genera that increased significantly (p <0.05, t-test, FDR corrected) in relative abundance in the after rain microbiome. As was observed with 16S rRNA amplicons in all samples (above), the urban signature bacteria *Arcobacter* increased by >50% in relative abundance following rain, though the increase was not statistically significant (Fig. 2B). *Legionella*, *Pseudomonas* and *Arcobacter* have all been previously associated with effluent contamination of urban waterways (20), supporting the significant role of increased effluent flow on the NSC microbiome. Increases in the relative abundance of other taxa such as *Francisella*, *Rickettsia* and *Chlamydia* that comprise pathogenic species (27, 28) and are usually not abundant in aquatic environments could be either a result of microbial influx from the effluent and/or the CSOs upstream. The decrease in the freshwater groups of Actinobacteria and Pelagibacteria after rain likely reflects a dilution effect on baseflow NSC waters from the increased effluent and CSOs flow. Several species, including *Francisella tularensis*, *Candidatus Nitrospira defluvii*, *Legionella longbeachae* and *Enterococcus faecalis*, were rare (<0.1% of the total sequences characterized by MyTaxa) in the before-rain microbiome but increased in relative abundance after rain to > 0.1% (Table S3). Most of these species are not common freshwater bacteria and are indicative of contamination.

**Figure 2:**
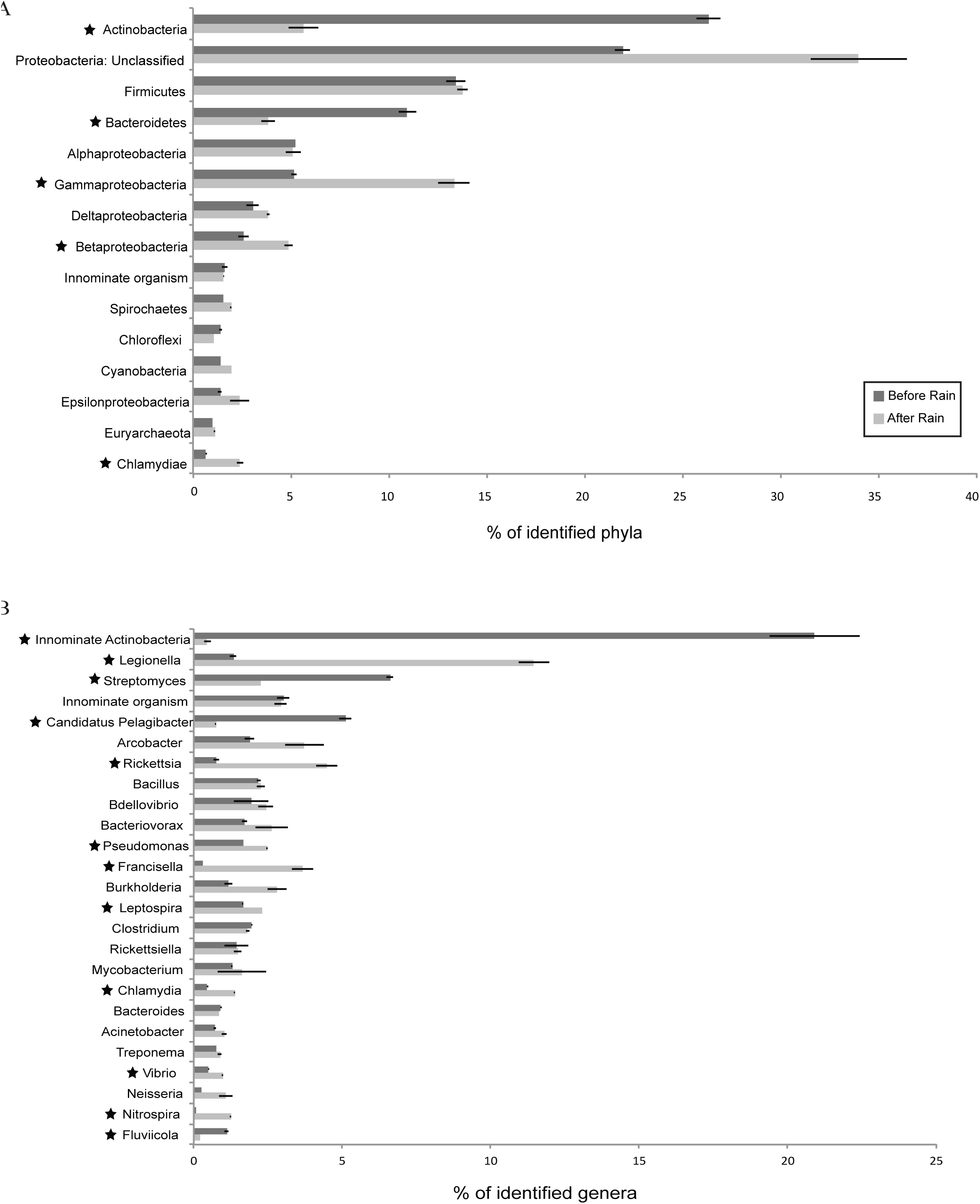
Rank-abundance plots for **(A)** phylum (Proteobacteria subdivided into classes) and **(B)** genus level classifications of metagenomic contigs from October 2013 before and after rain samples. The relative abundances of different taxa are averages of biological replicates for each sample (n=2). Based on taxon mean relative abundance across the samples, only the top 15 phyla and top 25 genera are shown. Phyla and genera highlighted with star symbol represent taxa with significant difference in relative abundance between the before and after rain microbiota (p < 0.05, t-test, false discovery rate corrected). ‘Innominate organism’ comprises contigs classified as organisms that either belonged to no known phylum/genus or a candidate phylum/genus.

### Population-level changes in response to rainfall in the North Shore Channel

We followed population-level trends for abundant organisms that exhibited large changes in their relative abundance after rain. Organisms most similar to *Legionella pneumophila* increased 10-fold in relative abundance after rain and also comprised the largest fraction of characterized species (11%) in the after-rain microbiome. Reads were recruited to the longest contig assigned to *L. pneumophila* in the rain-associated samples with roughly equal similarity (about 90-100% nucleotide identity) from each sample, suggesting the presence of the same population both before and after rain that increased substantially after rain (Fig. S4). This was supported by similarities in the average amino acid identity (AAI) of predicted protein coding genes from *L. pneumophila* before and after rainfall contigs (60% and 63%, respectively) to the genome sequences of the environmental isolate *L. pneumophila strain LPE509* and the clinical isolate *L. pneumophila subsp. pneumophila str. Philadelphia 1.* The AAI between genes attributed to *L. pneumophila* in the before and after rain metagenomes was 83%. Although genome pairs for the same species typically exhibit higher AAIs (~90%) (29, 30), 83% still signifies close genetic relatedness and not necessarily distinct populations. Overall, these results indicate that the before and after rain *Legionella* are members of the same species, but different from any currently known, sequenced members of *Legionella.* The discordance between our *Legionella*-like organisms and well-characterized *L. pneumophila* strains also makes it unclear if the corresponding populations are pathogenic, although a few predicted genes (1 and 3 for the before and after rain metagenomes, respectively) had high identity matches (>90%) to known *L. pneumophila* virulence genes in the virulence factor database (http://www.mgc.ac.cn/VFs/). Organisms within *Legionella* have been associated with artificial aquatic environments such as water distribution systems and cooling towers in buildings (31, 32) as well as WWTP effluent (20), thus their dramatic post-rain surge is not surprising.

Another notable increase in relative abundance after rain (~16-fold) was attributed to *Francisella tularensis*, an organism with known soil- and water-borne pathogenic subspecies (26, 32). Using a similar approach as above, AAIs between genes attributed to *F. tularensis* in before and after rain samples and a reference genome of pathogenic subspecies *F. tularensis subsp. tularensis SCHUS4* were 47% and 54%, respectively. Similar AAI values were observed between the metagenomic sequences and genomes of low virulent subspecies of this organism.

The AAI between the before and after rain *F. tularensis* genes was 68%. Thus, sequences classified as *F. tularensis* in our samples likely share the same taxonomic order Thiotrichales, but are different species from the known *F. tularensis* and might represent different populations within the same genus in the before and after rain samples, although the low number of sequences in the before rain dataset could bias in AAI calculation.

We also evaluated the population dynamics for species that dramatically dropped in relative abundance after the rain. *Actinobacterium SCGC AAA027*-*L06* is a member of the ubiquitous freshwater *Actinobacteria* lineage acI-B (33), and the relative abundance of contigs affiliated with this organism decreased dramatically (43-fold) after rain. Read recruitment indicated similarity between the before and after rain populations, with reads from each sample sharing ~90-100% nucleotide identity to the largest contig of this organism, although fewer reads mapped to the contig from the after rain samples (Fig. S5). As with the *L. pneumophila* population, the 84% AAI between the before and after rain sequences indicates close genetic relatedness between the two populations. Furthermore, the AAIs with respect to the *Actinobacterium SCGC AAA027*-*L06* draft genome were similar for the sequences from the before and after rain microbial communities (81% and 83%, respectively), indicating close genetic relatedness to this organism. Members of the acI-B lineage have been detected in diverse freshwater habitats (20, 35, 36, 38) and tend to prefer oligotrophic environments due to their small cell-size and oligotrophic life strategies (19, 36). Their decrease in relative abundance after rain likely reflects the reduced influence of freshwater flow from Lake Michigan due to increased wastewater flow.

### Overall functional gene content in before and after rain microbial communities

Functional gene profiles revealed taxa-driven shifts in the microbial community functional potential after rain. Although many abundant Gene Ontology (GO) terms related to housekeeping functions, such as nucleic acid and small molecule binding, did not significantly change in relative abundance after rain (data not shown), we observed an increase of >50% of functions within the broad terms of transporter activity and carbohydrate metabolism after rain. These were primarily related to transmembrane and substrate-specific transporter activity and carbohydrate biosynthetic and metabolic processes, respectively (Fig. 3A). Genes related to multi-organism processes such as pathogenesis and conjugation were >50% more abundant after rain while the before-rain microbiome had >50% more functions related to catabolic process, amine metabolic process and phosphate containing compound metabolic process (Fig. 3A). Within the broad GOs, genes related to photosynthesis, biosynthesis of organic compounds such as amines, vitamins and pigments as well as the activity of enzyme groups oxidoreductase (acting on the CH-NH_2_ group of donors) and ligase (forming phosphoric ester bonds) were twice as abundant in the before-rain microbiome (Fig. S6).

**Figure 3:**
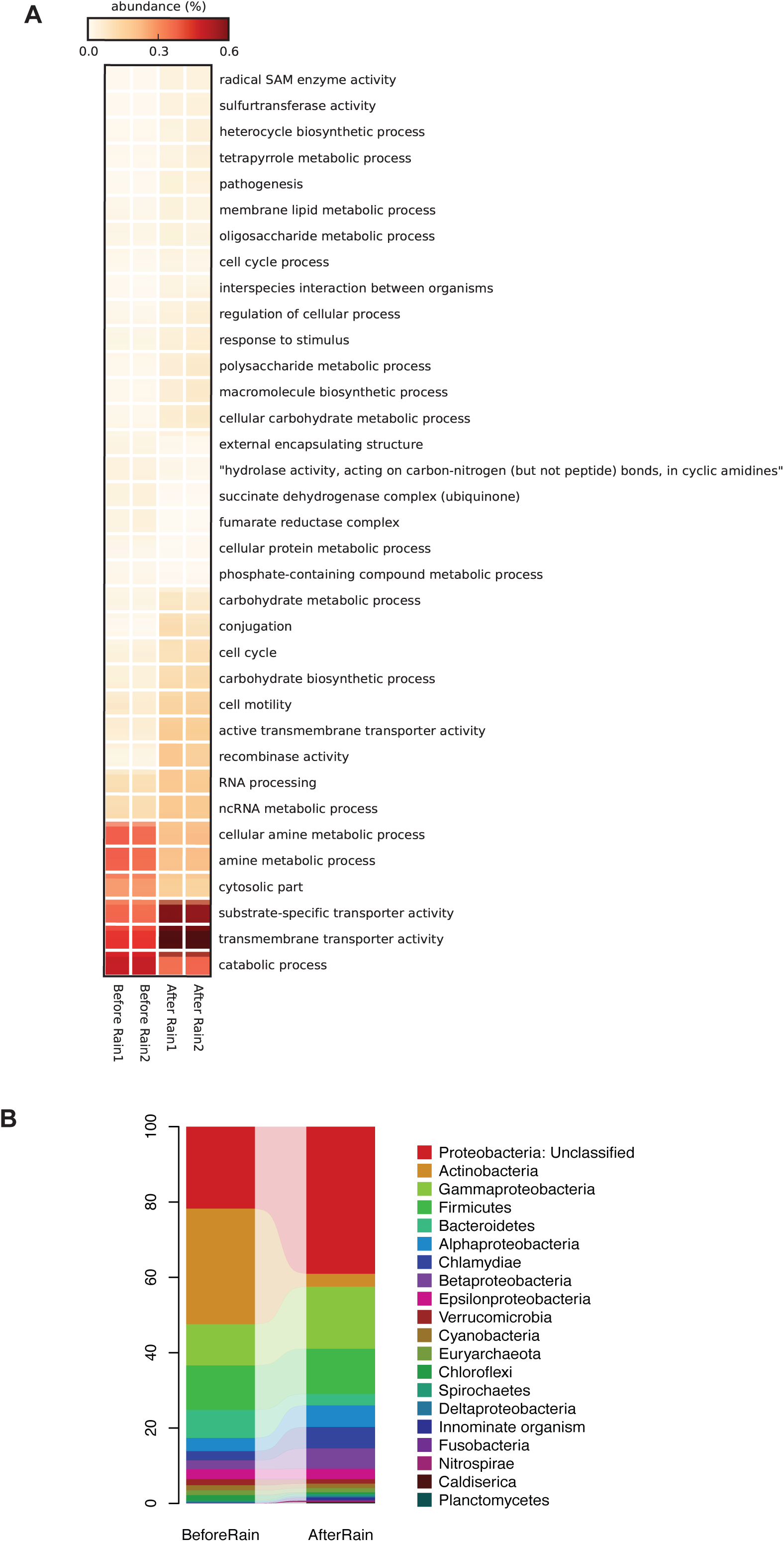
**(A)** Heatmap showing relative abundance (percentage of total predicted genes) at level 3 of Gene Ontology (GO) terms for the before and after rain microbiomes. GOs that had a higher relative abundance (> 50%) in one of the two groups (before/after rain) as compared to the other are shown. GOs that had less than 100 gene counts (*in situ* abundance) across all the samples have been excluded from the plot. Samples numbered 1 and 2 for each time point represent biological replicates. **(B)** Taxonomic composition at phylum level of genes from the rain-event microbial communities classified within the GO term ‘transmembrane transporter activity’. Relative abundances are a fraction of total sequences identified at phylum level.

Within the broad GO term of transporter activity, genes related to substrate-specific transmembrane transporter activity, specifically organic acid and ion transmembrane transporter activity, doubled in relative abundance after rain from an average of 0.06% to an average of 0.12% (Fig. S6). Genes encoding all transmembrane transporters were primarily attributed to Actinobacteria (31% of the identified sequences at phylum level) and unclassified Proteobacteria (22%) before rain, whereas unclassified Proteobacteria (39%) and Gammaproteobacteria (16%) were the major groups encoding transporters after rain (Fig. 3B). Gammaproteobacteria harboring transporter genes increased by 51% after rain while Actinobacteria encoding these genes exhibited more than 9-fold decrease, mirroring the shifts observed for the overall taxonomic profiles for these groups (Fig. 2, 3B). Genera contributing to the increase in Gammaproteobacterial sequences included *Legionella*, *Francisella* and *Pseudomonas*, exhibiting a pattern similar to the shifts in their relative abundance in the overall microbial community. Furthermore, as with the overall microbial community, *Actinobacterium SCGC AAA027*-*L06* (unclassified at genus level) contributed the largest fraction of sequences encoding transmembrane transporter activity genes within Actinobacteria in the before rain community. Interestingly, based on the functional gene content of organisms with dominant shifts in their relative abundance, those organisms that increased after rain had a higher proportion of their genes affiliated to transporter functions compared to those that dropped in abundance after rain. For instance, 3.7% and 6.8% of the *L. pneumophila* and *F. tularensis* genes, respectively, were associated with transmembrane transport,whereas *Actinobacterium SCGC AAA027*-*L06* and the genus *Pelagibacter* had ≤ 2%. Thus, the increase in transporter functions following the rain appears to be directly associated with an increase in the relative proportion of a subset of the organisms that harbor these functions rather than an increase in the distribution of these genes across the community. Organisms with transmembrane transporter genes, especially for organic substrates like organic acids, may be more suited to take advantage of the heterogeneous environment resulting from stormflow conditions.

Further evidence that changes in community composition drove the overall changes in the metabolic capacity came from genes that decreased in relative abundance after rain, such as those encoding biosynthesis of organic substances, which mirrored the overall shifts in taxa (Fig. 2); Actinobacteria (39% of the identified sequences at phylum level) and unclassified Proteobacteria (31%) were the major taxa encoding organic substance biosynthesis before rain and unclassified Proteobacteria (45%) and Gammaproteobacteria (13%) after rain. The shortterm nature and lack of gene expression data makes it difficult to know about the viability and activity of these organisms, but taxa-driven shifts in community functional potential were recently observed in another river in response to sewage and terrestrial-derived organisms (15).

### Biodegradation and antibiotic resistance gene abundance before and after rain

In addition to the GO-based functional analysis, we examined how rainfall impacted biodegradation and antibiotic resistance gene content. Predicted ORFs from both the before and after rain metagenomes were searched against a compiled database of protein sequences of microbial enzymes involved in the degradation of 12 different compounds associated with wastewater contamination, stormwater runoff, and WWTP effluent input (Fig. 4A). We detected biodegradation genes (BDGs) in both the before and after rain samples for 8 out of the 12 contaminants tested, but observed a significant increase (p < 0.05, t-test) in the relative abundance of genes involved in the degradation of nicotine, phenol, 1,4-dichlorobenzene and pentachlorophenol and a decrease (p < 0.05) in cholesterol degrading genes after rain (Fig. 4A). Additionally, the total relative abundance of all BDGs was significantly higher in the after rain sample (p < 0.05, t-test). BDGs before rain were primarily affiliated with unclassified Proteobacteria and Actinobacteria (35% and 30% of the identified sequences at phylum level, respectively), with the profile shifting to unclassified Proteobacteria and Betaproteobacteria (49% and 19%, respectively) as the dominant members of the community after rain, similar to the overall taxonomic shifts described above. These results reflect the increase in effluent flow from the WWTP as well as the suspected presence of these compounds in untreated wastewater and CSOs (3,38,40,42) (Fig. 4A).

**Figure 4:**
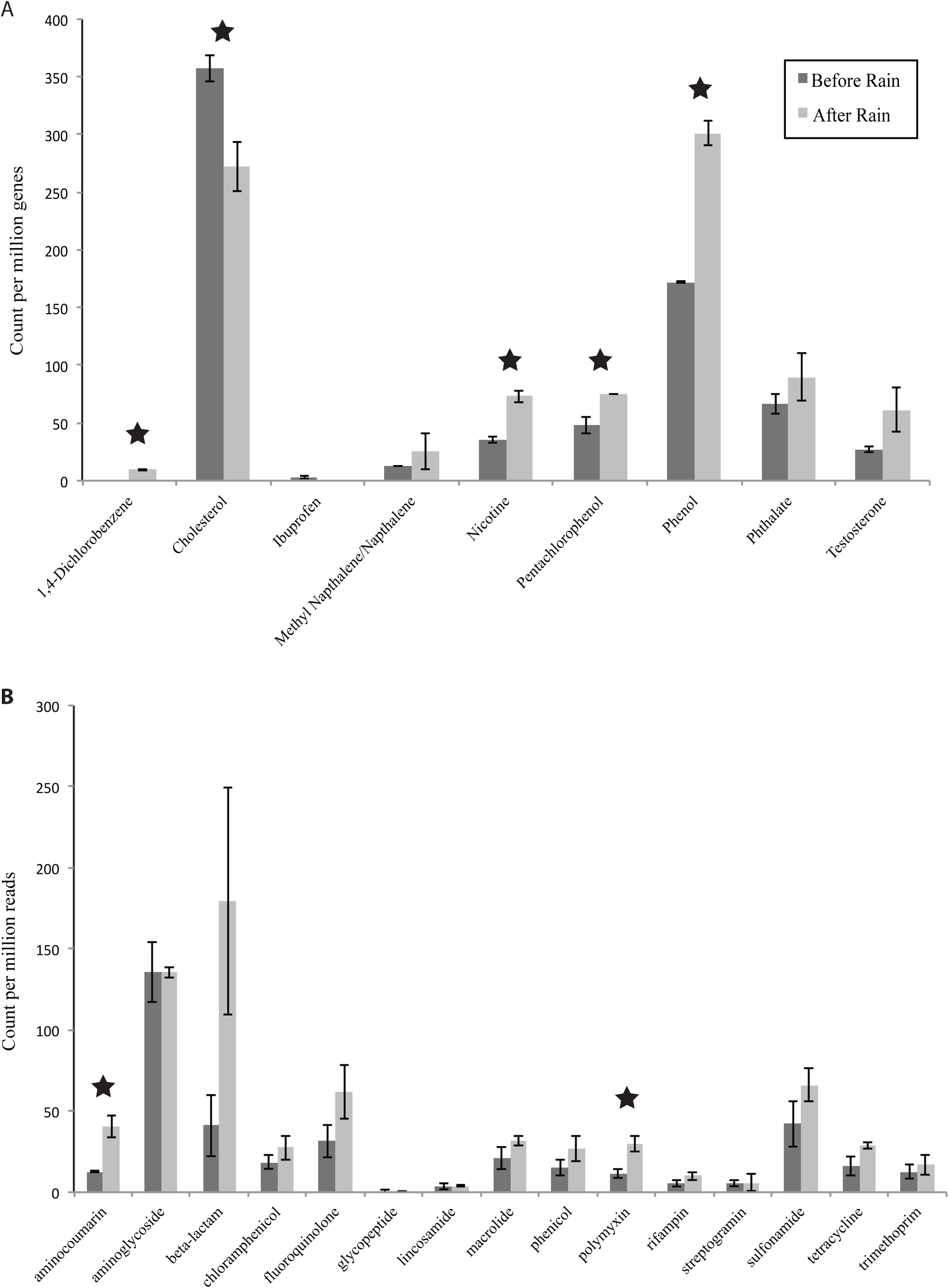
Relative abundance of **(A)** biodegradation genes (BDGs) and **(B)** antibiotic resistance genes (ARGs) in the before and after rain microbial communities. Relative abundance of BDGs refers to gene count (*in situ* abundance) per million genes per library averaged for each sample for their replicates (n=2) (see Methods section). For ARGs, relative abundance refers to read count per million reads per library averaged for each sample for their replicates. BDGs and ARGs with significant difference in relative abundances between the two time points (p < 0.05, t-test) are highlighted with stars.

Changes in the relative abundance of antibiotic resistance genes (ARGs) after rain were evaluated using the Comprehensive Antibiotic Resistance gene Database (CARD). As only a few ORFs (~10 per library) could be classified as ARGs from both the time points, we queried the unassembled paired-end reads against CARD. This resulted in several hits for various ARG categories in both time points (0.04% and 0.07% of the total number of reads for before and after rain samples, respectively) and revealed notable increases in the relative abundance of several ARG classes after rain (Fig. 4B), including significant increases in aminocoumarin and polymyxin resistance genes (p < 0.05, t-test). As with the BDGs, the total relative abundance for all ARGs pooled together for each time point was significantly higher in the after rain sample (p < 0.05, t-test). Increases in ARGs with urban-impacted stormflow was recently observed elsewhere as well (14), indicating that this could be a significant and underexplored effect of stormflow. Reads with high matches to ARGs were queried against metagenomic contigs, revealing that unclassified Proteobacteria and Firmicutes were the abundant ARG-carrying phyla (40% and 23% of the identified sequences at phylum level, respectively) in the before rain microbiome whereas unclassified Proteobacteria (50%) and Gammaproteobacteria (24%) were the dominant groups after the rain. This further supports the importance of taxa driven changes on gene content.

The results for both community composition and functional gene analysis provide evidence for the significant influence of WWTP effluent input on the microbial community, particularly from increased effluent flow-rates associated with heavy rain. Overall, this study revealed a shift in microbial community composition following rain from organisms frequently associated with freshwater systems towards organisms associated with urban impacted waters (9, 20, 21) as well as a shift in functional gene content. The increased relative abundance (and possibly actual abundance) of BDGs and ARGs along with the increase in genes associated with conjugation and pathogenesis in the after rain microbiome highlight the environmental and public health implications of stormflow in urban waterways. The extent to which these changes in gene content are expressed metabolically and persist is unknown. Although the WGS metagenomic analysis of a single rainfall event limits the scope of interpretations that can be drawn, our results provide substantial insights into microbial community dynamics in an urban stream during stormflow conditions, highlighting the need to investigate the urban stream microbiome with longer temporal scales and systematic sampling design to better predict the impact of rain associated stormflow events.

## Materials and Methods

### Site description and sample collection

The North Shore Channel (NSC) is a 12.3 km long man-made stream of the Chicago Area Waterway System that receives input from the O’Brien Water Reclamation Plant, a WWTP that serves over 1.3 million people residing in a 365 km^2^ area and releases effluent into the NSC (http://www.mwrd.org/irj/portal/anonymous/waterreclamation). Our study site is approximately 1 km downstream of the WWTP outfall (Fig. S1). The NSC also has 48 CSOs along its course, six of which are located within about 1 km upstream of WWTP, and two are located within 1 km downstream of the WWTP. These release excess stormwater mixed with untreated sewage into the river when the transport and storage capacity of the city’s sewage network is exceeded following high rainfall (http://www.mwrd.org/irj/portal/anonymous/overview)(Fig. S1). Water from the selected NSC site was sampled five times between 2013-2015 (0-1 m depth): three represent stream water during base flow (dry weather) conditions, and the other two represent stormflow (<24 hours after rainfall) conditions. We also sampled the WWTP effluent in October 2013 during baseflow conditions. Additional sample metadata and water chemistry are in Table S1.

Water was collected using a horizontal sampler (Wildco, Yulee, FL, USA) and passed on-site in succession through ~1.6 μm pore size glass fiber filters to remove larger particles (Whatman, Pittsburgh, PA, USA) and collected on a 0.22 μm pore size polycarbonate membrane filters (EMD Millipore, Billerica, MA, USA). WWTP effluent was collected from the WWTP outlet where the released effluent mixes with stream water. About 10L of water was filtered in duplicate for each sample and ~20 ml of the filtrate was transported back to lab for chemical analysis. Water Temperature, pH, conductivity and total dissolved solids were measured on-site using a portable water quality meter (Hanna Instruments, Woonsocket, RI, USA). Additional water chemistry analysis is described in Table S1.

### DNA extraction and sequencing

DNA was extracted from filters as described in (46). Briefly, filters were incubated in lysis buffer (50 mM Tris-HCl, 40 mM EDTA, and 0.75 M sucrose) containing 1 mg/ml lysozyme and 200 μg/ml RNase at 37 °C for 30 min. Subsequently, the samples were incubated with 1% SDS, 10 mg/ml proteinase K at 55 °C and rotated overnight. From the lysate, DNA was extracted using phenol:chloroform, followed by ethanol precipitation and elution in TE buffer.

Whole genome shotgun (WGS) metagenomic sequencing was done on the Illumina HiSeq (v1) with paired end format and read length of 150 bp at the Michigan State University Research Technology Support Facility. We obtained 2.82 and 3.18 Gbp of paired-end read data for the before and after rain samples, respectively. Replicate filters were sequenced at the University of Illinois at Chicago DNA Services Facility (DNAS) on a single lane of the Illumina HiSeq platform with paired end format and read length of 100 bp, yielding 4.04 and 1.31 Gbp of paired-end read data for the before and after rain libraries, respectively.

For 16S rRNA gene amplicon sequencing, 10-30 ng of DNA from each biological replicate (filter) were amplified with the V1-V3 primers 27F and 534R (47, 48). Amplicons were sequenced at the DNAS on the Illumina MiSeq platform with paired end format and read length of 300 bp. Between 28,933- 160,811 sequences per sample were obtained, with an average of 61,337 sequences per sample. All these sequence data have been submitted to the Sequence Read Archive at NCBI under accession number SRP080963.

### 16S rRNA gene based analysis of microbial community diversity

Paired-end barcoded reads of 16S rRNA gene amplicons were obtained for all the time points sampled and quality filtered using Trimmomatic (49) with a minimum average quality score of 20 across a 4-base sliding window and a minimum read length of 100 bp (including primer) post trimming. Trimmed, paired-end reads were merged using Pear (50), but owing to low yield of the merged reads, likely due to issues related to the MiSeq V2 kit chemistry, further analysis was only performed on the trimmed forward reads. Reads were analyzed using QIIME version 1.8.0 (51). Library statistics are summarized in Table S2. Chimeric sequences were removed using *identify*_*chimeric*_*seqs.py* with usearch61 denovo method and *filter*_*fasta.py.* Filtered sequences were clustered into operational taxonomic units (OTUs) at a 97% identity level using scripts *pick*_*otus.py* and *pick*_*rep*_*set.py* based on usearch61 denovo OTU picking. Representative OTUs were assigned taxonomy based on the Greengenes reference database (May 2013 version) using *assign*_*taxonomy.py* with uclust. OTUs occurring as singletons or with sequences from just one library were excluded from analyses. Community taxonomic composition and alpha diversity was performed using *summarize*_*taxa.py* and *alpha*_*diversity.py*, respectively, with a random subsample of 17,384 sequences per sample to avoid bias arising from variation in sequencing depth. Good’s coverage for each library was estimated using *alpha*_*diversity.py* and OTUs that included singletons, subsampled to an even depth of 18,289 sequences per library, the smallest library size.

### Metagenomic sequence assembly and phylogenetic classification

Raw metagenomic sequences were quality filtered using a Phred average per sliding window with quality threshold Q ≥ 20 and not allowing any N’s. Quality filtered coupled reads for each metagenomic library were assembled as described in (46). Coupled reads were first assembled into contigs with Velvet (52) and SOAPdenovo2 (53) separately, and input to Newbler 2.0 to obtain longer contigs with better N50 values (54). Additional metagenomic library statistics are provided in Table S3. Gene calling was done with MetaGeneMark (55). Due to uneven data yields from sequencing, we used assemblies from the first sequencing run for each sample as the representative sequences for annotations, and mapped the coupled reads from both the replicate libraries to these contigs for each sample to calculate the contig coverage in each library. The predicted protein coding genes for each dataset were used for phylogenetic classification of the corresponding contigs using MyTaxa (28) with a database of all sequenced bacterial and archaeal genomes (http://enve-omics.ce.gatech.edu/data/mytaxa) using DIAMOND blastp in the sensitive mode (56). Reads were mapped to contigs using blastn with cutoffs ≥ 50% alignment length, identity ≥ 97% and e-value ≤ 10^-10^. Contig coverage (sum of lengths of reads mapping to contig/contig length) was used as a proxy for *in situ* abundance in each library and calculated using the *BlastTab.seqdepth*_*nomedian.pl* script from the Enveomics bioinformatics toolbox (57). The script aai.rb from the same toolbox was used to calculate average amino acid identity (AAI) between any two sets of protein coding genes.

### Analysis of functional gene content and antibiotic resistance genes

Predicted metagenomic genes were searched against the SwissProt database (58) using blastp and cutoffs of at least 40% sequence identity, 70% coverage of the query sequence and e-value ≤ 10^-10^. The SwissProt match for the best hit for each query sequence was mapped to its corresponding Gene Ontology (GO) term (59), followed by binning the characterized genes at various depths (distance of a GO term from the parent node) of the GO database using in-house scripts. To evaluate the functional profile at a specific depth, *in situ* abundance for these GO terms was calculated using gene coverage (described above), and relative abundance for each GO term was obtained as a fraction of the total abundance of genes with identified functions in that library. The taxonomic affiliation of genes classified within a specific GO term was evaluated using MyTaxa, as described above.

To specifically evaluate the presence and abundance of genes involved in biodegradation of select wastewater contaminants in the rain-associated metagenomes, we created a database of protein sequences of enzymes related to degradation of select contaminants that are commonly found in WWTP effluent and sewage: testosterone; ibuprofen; caffeine; nicotine; cholesterol; 1,4-dichlorobenzene; methyl-naphthalene; pentachlorophenol; phenol; N,N, diethyl-3-toluamide; tetrachloroethylene and phthalate (3, 38–43). The enzymes were selected based on their role in the degradation pathways for these compounds (60), as well as the sequence availability in NCBI. This database is available from the corresponding author upon request. The predicted ORFs were searched against this database using blastp and the best hits were filtered at same thresholds used for SwissProt (above). Coverage estimates were used for calculating the *in situ* abundance for each BDG class, and normalized for each library by dividing the abundance of each BDG class by the total coverage of all predicted genes in that library and multiplying the result by 1 million to obtain gene count per million genes per library.

Antibiotic resistance genes in the rain-associated samples were identified by searching the predicted ORFs as well as paired-end metagenomic reads against the Comprehensive Antibiotic Resistance gene Database (CARD) (61) using blastp and blastx and a threshold of at least 80% sequence identity and 80% coverage of the query sequence (62, 63). Filtered reads for each library were binned into broad antibiotic resistance categories using the Resistance Gene Categories index file provided on CARD website (http://arpcard.mcmaster.ca/), and the read counts for each category were normalized for the library size as read count for ARG category per million reads per library.

### Microbial abundance estimation using fluorescence microscopy

October 2013 NSC samples were fixed with paraformaldehyde (1% final concentration) in triplicate and stored in 4°C. Samples were then vortexed and collected on 25 mm black polycarbonate filters (0.2 μm pore size) and stained with 5 μl of a 10 mg/ml DAPI (4’,6-diamidino-2-phenylindole) working solution diluted in 10X phosphate buffered saline (PBS). Microbial cells were enumerated (three slides from three replicate samples per time point) with an epifluorescence microscope (Zeiss Axio Scope.A1).

### Statistical analyses

Analysis of Similarity (ANOSIM) and Similarity Percentage (SIMPER) analysis on 16S and metagenomic community composition datasets, respectively, was performed using the R vegan package (64). The Statistical Analysis of Metagenomic Profiles (STAMP) software package was used for t-tests to evaluate differentially abundant taxonomic groups among the 16S rRNA gene and metagenomic datasets (65), and with R to evaluate differentially abundant physicochemical parameters, ARGs and BDGs. Principal Coordinate Analysis (PCoA, Bray Curtis metric) of OTUs (singletons removed and table subsampled to an even depth per sample) was performed with the Phyloseq package in R (66).

## Acknowledgements

This work was supported by the University of Illinois at Chicago. We thank Markeia Scruggs and Neil Mohindra for assistance with sampling and the personnel of the University of Illinois at Chicago DNA Services Facility for facilitating sample sequencing. We also thank the anonymous reviewers whose suggestions improved this manuscript.

## Supplemental Table and Figure Legends

**Table S1:** Water chemistry and environmental characteristics for North Shore Channel sampled time points.

**Table S2:** Sequencing statistics and diversity estimates for the 16S rRNA gene amplicon libraries used in the study.

**Table S3:** Sequencing statistics for the metagenomes used in the study.

**Table S4:** Rare species in before rain microbiome that were in the abundant fraction after rain.

**Fig. S1:** Map of the Chicago Area Waterway System (left panel) and the North Shore Channel (NSC) (right panel). Our study site at NSC is highlighted with an arrow. The point designated as WWTP on the right panel represents the O’Brien Water Reclamation Plant. Black dots along the stream represent locations for monitored CSO outfalls. CSO outfalls marked with red stars (locations A, B and C) recorded CSO events in the evening of Oct 5, 2013 with durations of 56, 50 and 5 minutes, respectively (http://www.mwrd.org/irj/portal/anonymous/overview).

**Fig. S2:** O’Brien Water Reclamation Plant effluent flow rate [million gallons per day (MGD)] and rain gauge data for the months of September and October 2013 (http://www.mwrd.org/irj/portal/anonymous/overview).The circled region of the plot corresponds to data around the rain event (10/5/2013), which is the focus of this study. No data was available for 9/17/2013 as the rain gauge was out of service.

**Fig. S3:** Community coverage estimates based on metagenomic reads generated using Nonpareil for the before and after rain metagenomes. Sample numbers 1 and 2 for each time point represent biological replicate libraries.

**Fig. S4:** Reads from before rain (top) and after rain (bottom) datasets were mapped to the longest contig attributed to *Legionella pneumophila* from the after rain metagenome. Reads for biological replicate libraries (n= 2) were pooled for both the before and after rain time points.

**Fig. S5:** Reads from before rain (top) and after rain (bottom) datasets were mapped to the longest contig attributed to *Actinobacterium SCGC AAA027*-*L06* from the before rain metagenome. Reads for biological replicate libraries (n= 2) were pooled for both the before and after rain time points.

**Fig. S6:** Heat map showing the relative abundance (percentage of total predicted genes) at level 4 depth of Gene Ontology (GO) terms for the before and after rain microbiomes. GO terms that had a higher relative abundance (> 100%) in one of the two groups (before/after rain) as compared to the other are shown, and terms that had less than a total of 75 gene counts across all the samples have been excluded from the plot. Samples numbered 1 and 2 for each time point represent biological replicates.

